# Bacteriophage in combination with ciprofloxacin against *Pseudomonas aeruginosa* infections in diabetic foot ulcer patients

**DOI:** 10.1101/2023.10.03.560795

**Authors:** Sha Liu, Shenoi Goonetilleke, Karen Hon, Isabella Amy Burdon, George Spyro Bouras, Neil McMillan, Alkis J Psaltis, Peter-John Wormald, Robert Fitridge, Sarah Vreugde

## Abstract

**Background:** In diabetic foot ulcer (DFU) patients, *Pseudomonas aeruginosa* (*P. aeruginosa*) infections are linked to poor wound healing. The ineffectiveness of antibiotics against these infections promotes the emergence of multidrug-resistant (MDR) strains. Bacteriophage (phage) therapy has recently gained popularity as an alternative to antibiotics.

**Methodology:** Bacterial and viral swabs and tissue were obtained from DFU infections (DFI). Bacteria were cultured followed by MALDI-TOF MS for identification. 16S rRNA long-read sequencing was used to identify the microbiota. Bacteriophages were isolated and underwent transmission electron microscopy, genomic sequencing, and stability testing. The antimicrobial activity of phages alone and in combination with ciprofloxacin against *P. aeruginosa* planktonic cells and biofilm grown *in vitro* and in *ex vivo* tissue was tested by measuring the optical density (OD), crystal violet assays and live/dead staining with visualisation using confocal scanning laser microscopy respectively.

**Results:** A total of 34 DFI patients were recruited from which microbiota were analysed for 25 patients. *P. aeruginosa* was the most prevalent pathogen cultured and was one of the top 6 most prevalent and abundant species in the microbiota analysis. Phage APTC-PA18 was isolated from DFIs, belonged to the *myoviridae* family and was strictly lytic. PA18 was stable between 4 and 70 degrees Celsius and between pH 3 and 11. Seven of eight *P. aeruginosa* clinical isolates were sensitive to APTC-PA18, and when APTC-PA18 was combined with ciprofloxacin against planktonic and biofilm of *P. aeruginosa*, synergistic effects were observed *in vitro* and in DFI tissue samples.

**Conclusion:** Phage APTC-PA18, when combined with ciprofloxacin, has the ability to kill *P. aeruginosa* clinical isolates both *in vitro* and *ex vivo* and is a promising treatment option for *P. aeruginosa* infections in DFUs.

## 1. Introduction

*P. aeruginosa* is a gram-negative opportunistic pathogen that can cause a number of nosocomial infections and is responsible for complex recurrent infections in immunosuppressed hosts ^1, 2^. Diabetes-related foot diseases (DFDs) are among the top ten causes of hospitalisations in Australia with an annual cost of $1.6 billion ^3^. *P. aeruginosa* is the most prevalent gram-negative pathogen across all types of infections in the DFUs ^2, 4^. *P. aeruginosa* DFU infections (DFI) can result in serious tissue damage that associated with poor prognosis, which may eventually necessitate amputation of a limb ^5^. *P. aeruginosa* DFIs are notoriously difficult to treat, especially due to limited antibiotic susceptibility. Resistance to broad-spectrum antibiotic treatments is achieved through the utilisation of intrinsic mechanisms including low permeability of the *P. aeruginosa* outer membrane, biofilm formation and its ability to acquire and express multiple resistance mechanisms including porin deletions and overexpression of efflux pumps^6, 7^. Therefore, the rapid emergence of multi-drug resistant (MDR) *P. aeruginosa* strains and the rise in the number of associated infections globally is a concerning trend that calls for investigation into alternative therapies for this serious public health problem.

Bacteriophage (phage) therapy is a treatment option that can potentially be used in combination with antibiotics to treat MDR *P. aeruginosa* infections ^8, 9^. Some advantages of phage therapy over antibiotic therapy include increased specificity to their hosts, which reduces the likelihood of secondary infections, and a more targeted treatment because phages replicate at the site of infection ^8^. Preclinical trials have shown that phage is effective in killing planktonic *P. aeruginosa* and penetrating and disrupting *P. aeruginosa* biofilms ^9^. Phage therapy used in combination with traditional antibiotic treatment has shown synergistic reductions in *P. aeruginosa* growth and eradication of biofilms both *in vitro* and *in vivo* ^7, 10^. Despite the effectiveness of phage combination treatments against planktonic and biofilm *P. aeruginosa*, the use of phage therapy has been limited and novel phages that can be used to treat such infections are urgently needed. The purpose of our study was to investigate alternative *P. aeruginosa* infection management strategies for DFUs.

## 2. Materials and Methods

### 2.1: Human Ethics approval and participant recruitment

The Central Adelaide Local Health Network Human Research Ethics Committee (HREC/14/TQEH/603) approved the collection of swabs and tissue from patients attending the Multidisciplinary Foot (MDF) clinic at The Queen Elizabeth Hospital (TQEH) for this study. Written informed consent was obtained from participants before collection. DFUs were determined to be clinically infected as per the International Working Group on the Diabetic Foot (IWGDF) and The Society for Vascular Surgery Wound, Ischaemia, foot Infection (WIfI) ^11, 12^. Patients excluded from recruitment were those unable to give consent and those under the age of 16. Demographic data and medical history were recorded for all participants.

### 2.2: Sample collection

The Levine method of swabbing was used in the collection of bacterial swabs and microbiome swabs ^13^. Swabs were collected from DFIs after the wound had been cleansed, using sterile saline and gauze, and debrided as per the clinical therapeutic plan. Viral swabs for phage isolation were taken from DFIs in the same method. The swab was then suspended in a 4 ml tryptic soy broth (TSB) (Oxoid, Thebarton, SA, Australia), transported to the lab and stored at 4°C for up to 1 week.

### 2.3: Bacterial strain isolation and identification

100 μL of serially diluted bacterial suspension was pipetted onto sheep blood agar plates (Thermo-fisher Scientific, Thebarton, SA, Australia). The plate was then incubated at 37°C supplied with 5% extra CO_2_ (Panasonic Healthcare Co., Tokyo, Japan) overnight. The following day, individual colonies with distinct morphologies were sent for identification via matrix-assisted laser desorption ionization-time of flight mass spectrometry (MALDI-TOF MS) (Bruker, VIC, Australia). Clinical isolates were stored in 25% glycerol in TSB at -80°C.

### 2.4: Microbiome DNA extraction

PureLink™ Microbiome DNA Purification Kit (Thermo Fisher Scientific, Waltham, MA, USA) was used to extract microbiome DNA according to the manufacturer’s instructions. DNA was then quantified using Nanodrop 2000 (Thermo fisher scientific, Waltham, MA, USA).

### 2.5: Barcoding DNA and Library Preparation

50 ng DNA was mixed with Rapid Barcodes (Oxford Nanopore Technologies; Oxford; UK) and incubated at 30°C for 2 minutes and then at 80°C for 2 minutes. 25 samples were pooled in equal amounts and the library was prepared according to the manufacturer’s instructions.

### 2.6: Priming and loading the SpotON flow cell

The flow cell (R9.4.1; Oxford Nanopore Technologies; Oxford; UK) priming mix was prepared and primed according to the manufacturer’s instructions. This was followed by adding 75 μL of sample via the SpotON sample port.

### 2.7: 16S rRNA sequencing result analysis

The 16S rRNA data was generated on a MinION platform (Oxford Nanopore Technologies, Oxford UK). Basecalling was conducted using the FAST configuration with Guppy version 6.1.5. The full pipeline used to analyse the resulting fastq files was https://github.com/gbouras13/16S_Emu_Pipeline and was created using Snakemake ^14^. Fastq files were filtered using filtlong, keeping all reads with a minimum quality score of 8 and a length between 1300 and 1700 bp. Nanoplot was used to generate quality control plots on the filtered fastq files ^15^. Species level relative abundances were determined using Emu v 3.3.1. Krona was used to visualise the relative abundances for each sample ^16^.

### 2.8: Minimum inhibitory concentration (MIC) determination of *P. aeruginosa* DFUs clinical isolates

MIC assay was used to determine the antibiotic resistance profile of *P. aeruginosa* CIs ^17^. The antibiotic tested was ciprofloxacin, obtained from Sigma-Aldrich (Castle Hill, NSW, Australia). The clinical isolates were deemed susceptible, intermediate, or resistant as per the Clinical and Laboratory Standards Institute (CLSI) recommendations (Adelaide Pathology Partners, Mile End, VIC, Australia)^18^.

### 2.9: Phage isolation from DFUs viral swabs

*P. aeruginosa* phage was isolated from viral swabs that were harvested from patients with *P. aeruginosa* positive. Phage was isolated using the double-layer agar method, as described previously^19^. Individual plaques with different morphologies were picked using pipette tips for propagation. After this processing, the phage was amplified and used the double-layer spot assay (DLSA) as described previously^19^.

### 2.10: Phage DNA extraction and genomic analysis

A phage DNA isolation kit (Norgen Biotek Corp, Thorold, ON, Canada) was used to carry out DNA extraction in accordance with the protocol provided by the manufacturer. Paired-end sequence reads were obtained on an Illumina sequencer (SA Pathology, SA, Australia). Reads were trimmed and filtered for quality using fastp ^20^. The genome assembly was performed using Unicycler v 0.5.0. The sequences were verified using QUAST ^21^ and annotated with Pharokka v 1.1.0 ^22^. Genomic analysis was conducted with the non-redundant (nr) NCBI database using the web-interface of MEGABLAST ^23^.

### 2.11: TEM imaging of phages

Transmission electron microscopy was carried out and electron micrographs of the phage were obtained and analysed. A modified protocol ^24^ was used. 5 μL of phage was placed on the coated side of a carbon/formvar grid (ProSciTech Pty Ltd., Kirwan, QLD, Australia) for 3 min before being wicked dry with filter paper. 5 μL of electron microscopy fixative (1.25% glutaraldehyde, 4% paraformaldehyde in PBS to which 4% sucrose had been added) was then placed on the grid for 2 minutes, followed by 5 μL of 2% uranyl acetate for 2 min. The samples were examined in an FEI Tecnai G2 Spirit 120 kV TEM (FEI Technologies Inc., Hillsboro, OR, USA).

### 2.12: Thermal and pH stability tests

Thermal stability tests were conducted by adding 100 μL phage working stock (10^10^ PFU/mL) that was resuspended in SM buffer and incubating for 1 hour in an Eppendorf Thermomixer Compact (Sigma-Aldrich, Castle Hill, NSW, Australia) at 4°C, 30°C, 37°C, 40°C, 50°C, 60°C, 70°C and 80°C.

PH stability tests were carried out on the phage, as described previously ^25^. 10 μL of each phage (10^9^ PFU) was suspended in 90 μL of the pH-adjusted SM Buffer (pH 3 to 12) and left to incubate at room temperature for 1 hour. The pH and thermal stability of the phage were determined by measuring the phage titration.

### 2.13: Inhibition Assay

PAO1 was infected with phage at MOI = 0, 0.1 and 1. In brief, overnight cultures were prepared and the OD_600_ was adjusted to 0.2. The required volume of phage stock solution (10^8^ PFU/mL) was added to the corresponding suspension to achieve an MOI = 0, 0.1 and 1. The samples were incubated at 37°C with 180 rpm shaking. Every 30 min for up to 6.5 hours, the OD_600_ value was measured. The experiment was repeated 3 times.

### 2.14: Host range testing

100 μL of an overnight culture of each CI was added to molten 0.4% TSA and plated on 1.5% TSA plates. 5 μL droplets of each dilution of phage were added in triplicate and left to dry. Then the plate was incubated at 37°C overnight. Sensitive CIs were identified by the formation of clear plaques or lysis, semi-sensitivity was identified by the formation of semi-opaque plaques or partial lysis and non-sensitive CIs by no plaque formation or no lysis ^26^.

### 2.15: Effect of phage in combination with antibiotics on planktonic *P. aeruginosa*

5 x10^5^ CFU/mL of each CI were added into 96-well microtiter plates. Treatments included phage(MOI=1), ciprofloxacin (½ MIC) or a combination of both. An untreated and TSB was used as a positive and negative control. Plates were incubated overnight at 37°C and the optical density (OD) readings at 595nm were recorded using an iMark™ Microplate Absorbance Reader (Bio-Rad laboratories, Hercules, CA, USA). Synergistic killing of planktonic cells is defined as a greater reduction in viability than can be expected when adding the effects of individual treatments ^27^.

### 2.16: Efficacy of phage combined with ciprofloxacin on *P. aeruginosa* biofilms

A 1.0 MCF *P. aeruginosa* culture suspended in 1:15 dilution in TSB was added 180 μL per well to a 96-well U-shaped microtiter plate (Thermo-Fisher, Roskilde, Denmark). Plates were incubated for 24 hours to allow biofilm formation. Biofilms were then treated with 10^8^ PFU/mL of phage and 1MIC, 2MIC and 3MIC of ciprofloxacin for 24 hours at 37°C on a rotating platform at 70 rpm. 24 hours post-treatment, wells were washed twice and stained with 180 μL per well of 0.1% crystal violet (Sigma-Aldrich, Castle Hill, NSW, Australia) for 15 minutes and 180 μL per well of acetic acid was added to the plates to elute the crystal violet stain. The OD readings at 595 nm were recorded using an iMark™ Microplate Absorbance Reader (Bio-Rad laboratories, Hercules, CA, USA). A synergistic reduction in biofilm biomass is defined as a greater reduction in biofilm biomass than can be expected when adding the effects of individual treatments ^27^.

### 2.17: Phage-antibiotic combination treatment against *P. aeruginosa* biofilm DFU tissue

Patients with confirmed *P. aeruginosa* infection were selected for this experiment. Briefly, tissue from the infection site was harvested after being rinsed with sterilized saline. The fresh tissue was placed into Dulbecco’s Modified Eagle’s Medium (Life Technologies, Carlsbad, California, USA) on ice and divided into 4 equal pieces and treated with PA18 phage (10^8^ PFU), 3 MIC ciprofloxacin, PA18 phage (10^8^ PFU) in combination with 3 MIC ciprofloxacin or left untreated. After 24 hours of treatment, tissues were placed on the slides and stained with a LIVE/DEAD BacLight Bacterial Viability Kit (Life Technologies Australia, Mulgrave, Victoria, Australia) according to a published protocol^26^. The stained tissue was then examined at 10X magnification using a confocal laser scanning microscope (Zeiss LSM700, Carl Zeiss AG, Oberkochen, Germany).

### 2.18: Statistical analyses

GraphPad Prism 9 software (GraphPad Prism version 9.0.0 for Windows, GraphPad Software, San Diego, CA, USA) was used for statistical analysis. Data were analysed using a one-way analysis of variance (ANOVA) followed by Tukey’s or Dunnett’s multiple comparison test. Significance was determined as p<0.05. All experiments were repeated three times and performed in triplicate or six replicates unless specified otherwise. The mean values of the three or six replicates were obtained with standard error of the means (SEM).

## 3. Results

### 3.1: *P. aeruginosa* is the most prevalent species cultured from DFIs

A total of n=34 DFU patients were included in this study. These included 25 males and 9 females. All patients had diabetes: n=19 patients were on insulin therapy and n=13 patients were on antibiotic therapy prior to being swabbed. From those 34 patients, a total of 45 CIs were isolated. These included 9 different genera and 19 different species of bacteria (Figure 1). The most prevalent species were *P. aeruginosa* (n=8), *Staphylococcus aureus* (n=6) and *Enterococcus faecalis* (n=5). The MIC of ciprofloxacin for each of the *P. aeruginosa* CIs was determined. Based on the CLSI breakpoints, *P. aeruginosa* APTC-DFI-1was deemed to be resistant to ciprofloxacin. Results are shown in Table 1.

**Figure 1.**
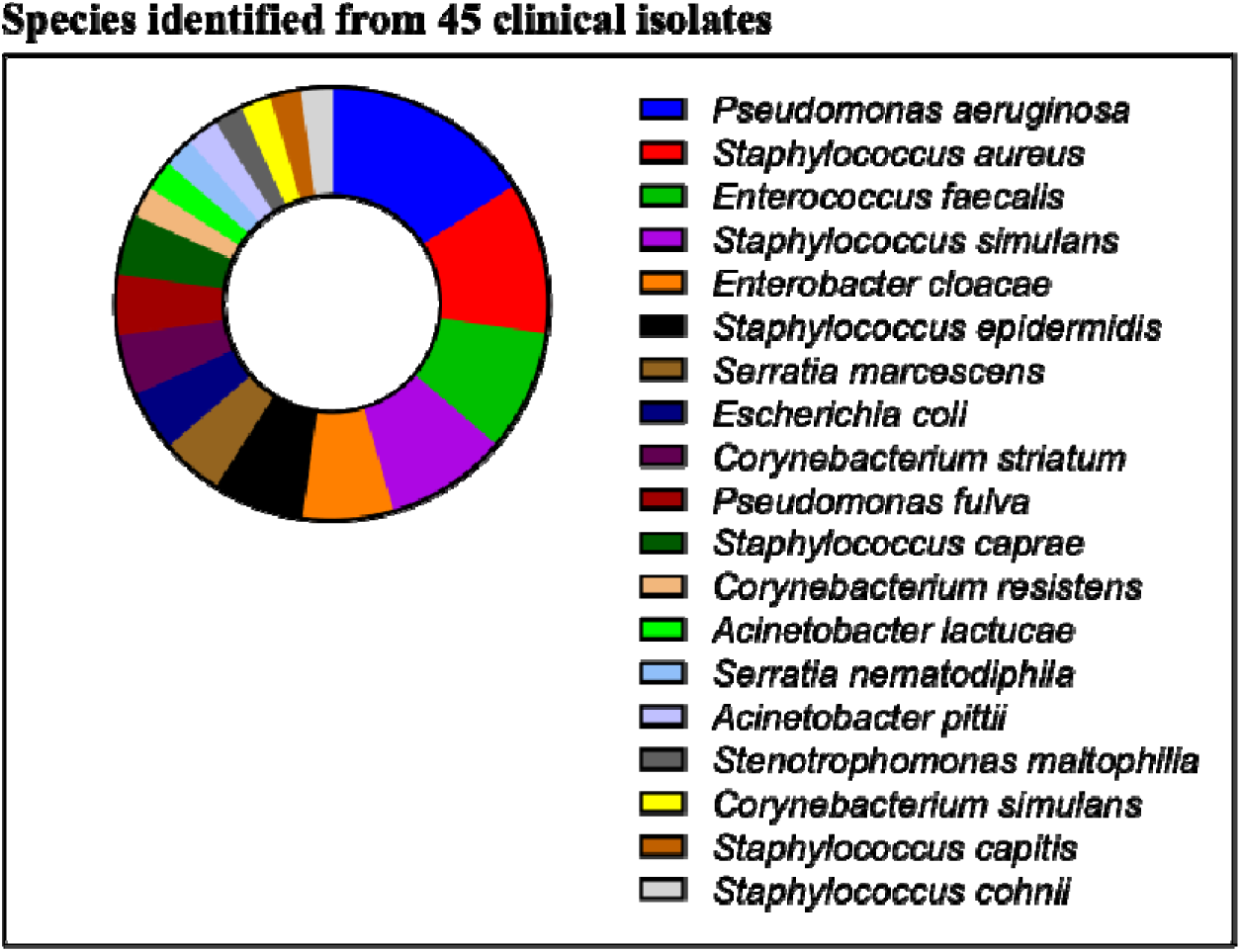
Bacteria isolated from DFUs. Different bacteria isolated from the DFIs of patients were identified and the proportion of each bacterium making up the n=45 CIs was listed in descending order. The CIs consisted of 9 different genera and 19 different species of bacteria.

**Table 1.** Minimum inhibitory concentrations (MIC) of ciprofloxacin for each *P. aeruginosa* CI from DFIs. Resistance to ciprofloxacin is denoted by “S” for susceptible, “I” for intermediate and “R” for resistant.

| DFI *P. aeruginosa* CI<br>Identification Number | Minimum inhibitory<br>concentration (MIC) of<br>Ciprofloxacin | Resistance to<br>Ciprofloxacin<br>(S=Susceptible)<br>(I=Intermediate)<br>(R=Resistant) |
| --- | --- | --- |
| APTC-DFI-1 | 16 | R |
| APTC-DFI-2 | 0.5 | S |
| APTC-DFI-3 | <0.125 | S |
| APTC-DFI-4 | <0.125 | S |
| APTC-DFI-5 | 0.25 | S |
| APTC-DFI-6 | 0.25 | S |
| APTC-DFI-7 | <0.125 | S |
| APTC-DFI-8 | 0.25 | S |

### 3.2: *P. aeruginosa* is one of the top two prevalent species in microbiota of DFIs

16S rRNA long-read sequencing was used to analyse the DFI microbiota from 25/34 patients recruited to this study. The bacterial genus for each patient and the top 10 most prevalent bacteria at the genus and species level are shown in Figure 2 A-C. The top 5 genera of bacteria in terms of mean relative abundance were *Staphylococcus*; *Finegoldia*; *Pseudomonas*; *Anaerococcus* and *Serratia*, all with mean relative abundance > 0.05. The top five species in terms of relative abundance were *Finegoldia magna*; *Staphylococcus epidermidis; Pseudomonas aeruginosa*; *Staphylococcus aureus* and *Enterococcus faecalis*.

**Figure 2:**
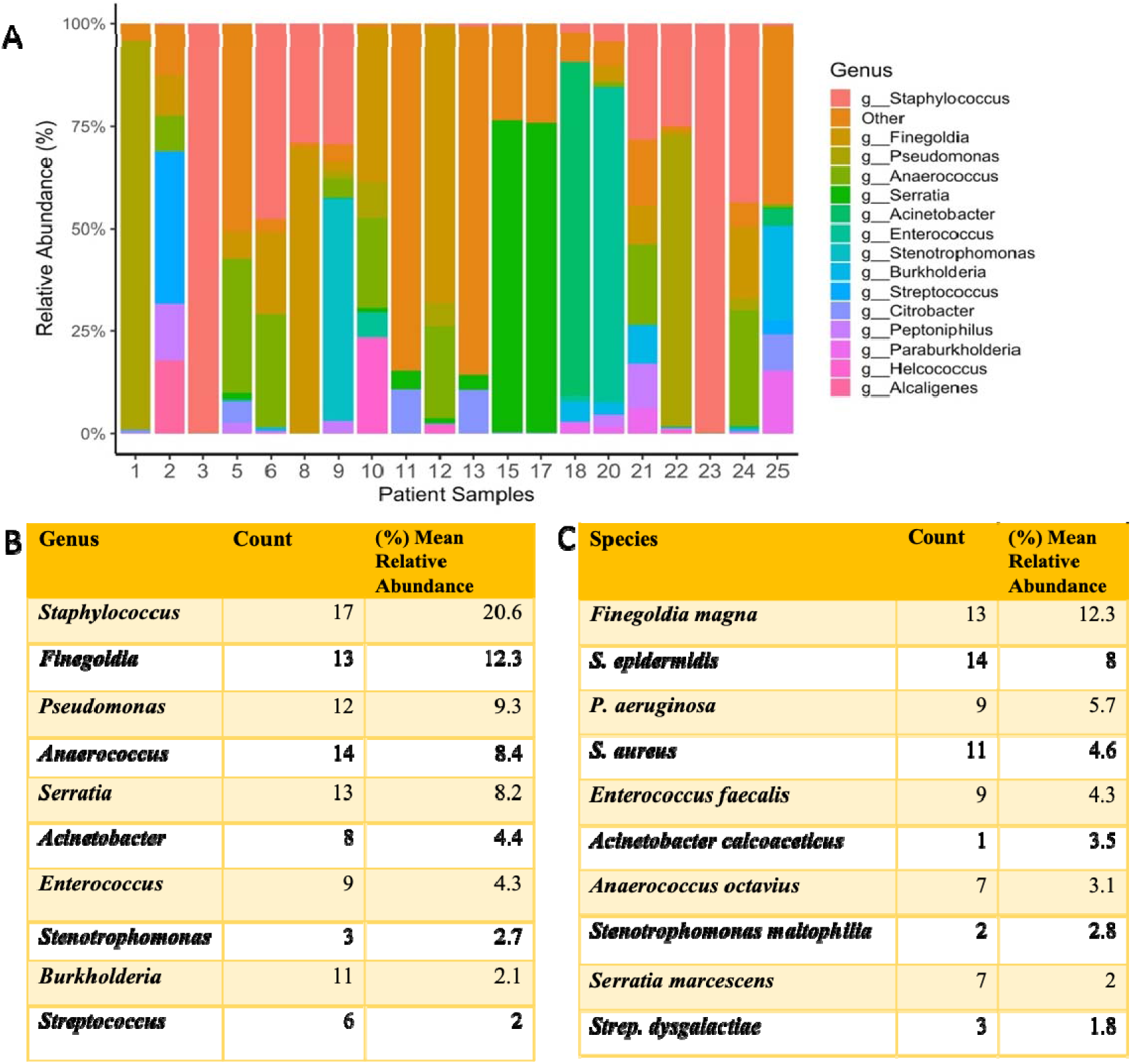
Microbiome 16s sequencing identified the prevalence of genus and species of pathogens. (A) the plot of 25 patients’ microbiome analysis; (B) Summary of top 10 prevalent bacterial genera; (C) Summary of top 10 prevalent bacterial species. Count=the number of samples that present the genus and species of pathogens. Mean=the average of the relative abundance of the genus and species of pathogens.

### 3.3: APTC-PA18 is a lytic *P. aeruginosa* phage isolated from DFI

One *P. aeruginosa* phage was isolated (designated as Adelaide Phage Therapy Centre *P. aeruginosa* phage 18 - APTC-PA18) from a DFI patient. Genomic sequencing determined that APTC-PA18 has a sequence length of 66459 bp. APTC-PA18 possessed genes encoding for endolysins and lacked genes associated with lysogeny, virulence, toxins, and antibiotic resistance (Figure 3 A) and was classified as lytic ^28^. MEGABLAST analysis determined that APTC-PA18 belongs to the family of *Myoviridae*. This was confirmed using TEM that indicated that APTC-PA18 had morphological characteristics consistent with that of the *Myoviridae* family including an icosahedral head (Figure 3 B).

**Figure 3:**
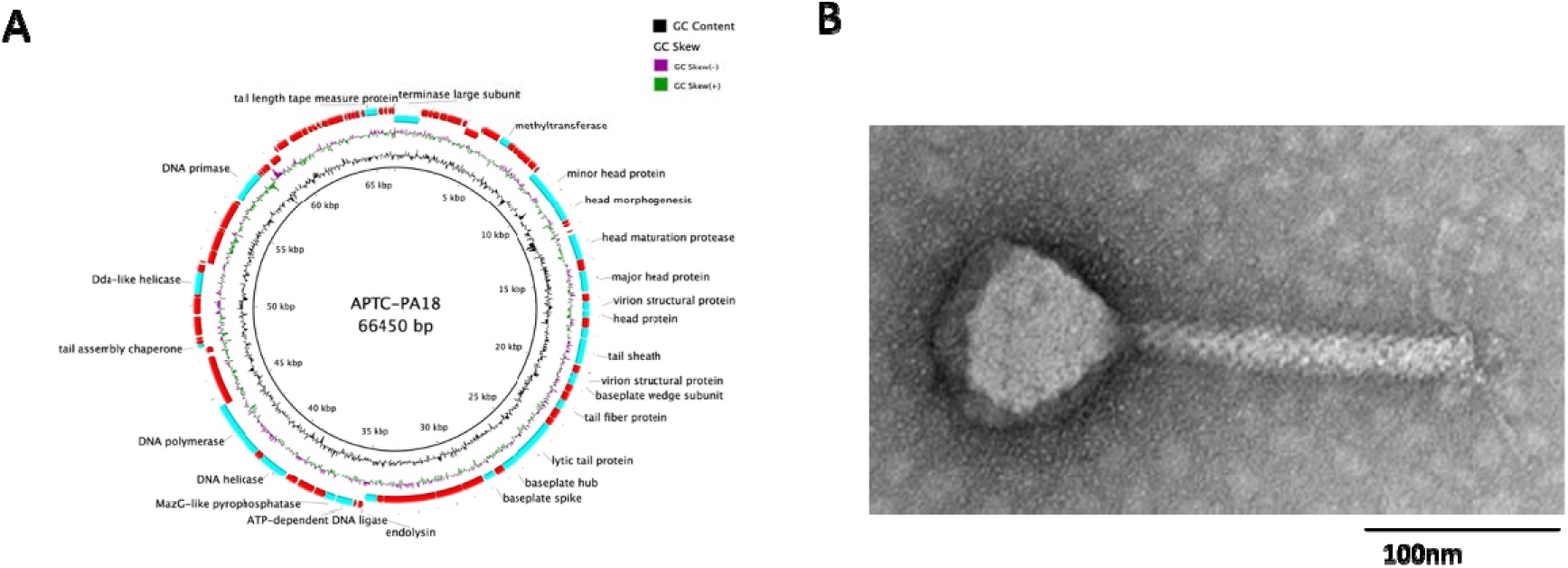
*P. aeruginosa* phage APTC-PA18 genomic characterization. (A) Phage APTC-PA18 genomic map. Circular representation of the genome of APTC-PA18. In blue are key features of the genomes and in red are hypothetical regions. (B) TEM of APTC-PA18 indicating phage belongs to the *Myovirida*e family with an icosahedral head. PA18=APTC-PA18.

### 3.4: Phage APTC-PA18 shows good stability and can kill *P. aeruginosa* at low concentrations

APTC-PA18 shows good stability at a pH range of 3-11 with no significant change or drop in phage viability when compared to phage viability at pH 7. At pH 12 no viable phage was detected (Figure 4 A). APTC-PA18 was stable at temperatures between 4°C – 60°C. At 70°C, there was a significant decrease in phage viability to 5.2 Log_10_ ^PFU/mL^ in comparison to 10.3 Log_10_ ^PFU/mL^ viability at 4°C. At 80°C, no viable phage was detected (Figure 4 B). Comparing the growth inhibition curves of APTC-PA18 relative to untreated control (MOI=0) showed that at MOI=1, a significant reduction in viable bacteria was observed at 150 min, whilst at MOI=0.1, a significant reduction in viable bacteria was obtained later, at around 210 min (Figure 4 C).

**Figure 4:**
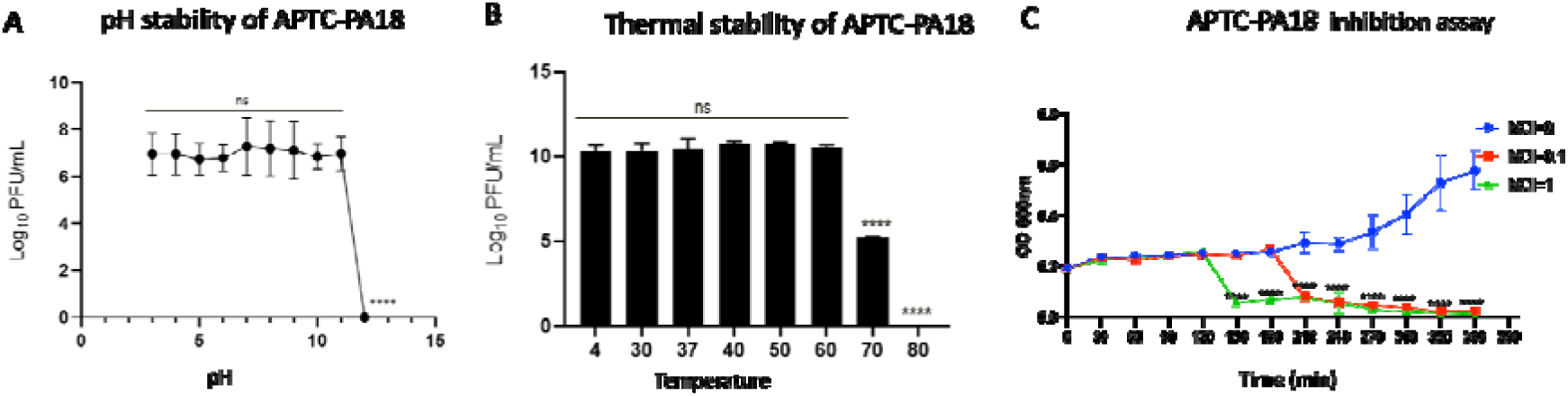
Phage APTC-PA18 characterization. (A) APTC-PA18 was incubated for 1 hour at RT at a pH range of 3-12; (B) PA18 was incubated for 1 hour at the temperatures 4°C, 30°C, 37°C, 40°C, 50°C, 60°C, 70°C and 80°C. Phage viability was indicated by Log_10_ ^PFU/mL^; (C) Bacteriophage APTC-PA18 inhibition assay. Inhibition of PAO1 reference strain after treatment with APTC-PA18 at a multiplicity of infection (MOI) = 0.1 (red) and 1 (green) compared to an untreated control (MOI =0) (blue). Data expressed as mean ± SEM for three independent experiments. Ns = not significant, ^****^ = p<0.0001.

### 3.5: APTC-PA18 can kill *P. aeruginosa* clinical isolates from DFIs

In order to determine the host range of PA18, phage sensitivity against all 8 *P. aeruginosa* CIs from DFIs was tested. APTC-PA18 exhibited a broad host range where 7 out 8 of the CIs were shown to be sensitive to APTC-PA18 and only 1 CI was shown to be non-sensitive (Table 2).

**Table 2.** Phage APTC-PA18 was tested against *P. aeruginosa* CIs from DFIs using a double agar spot assay. Sensitive CIs (+) formed clear plaques, semi-sensitive CIs (+/-) formed semi-opaque plaques and non-sensitive CIs (-) had no plaque formation on the bacterial lawn.

| DFI *P. aeruginosa* CI Identification Number | Sensitivity to bacteriophage APTC-PA18 |
| --- | --- |
| APTC-DFI-1 | + |
| APTC-DFI-2 | + |
| APTC-DFI-3 | + |
| APTC-DFI-4 | + |
| APTC-DFI-5 | - |
| APTC-DFI-6 | + |
| APTC-DFI-7 | + |
| APTC-DFI-8 | + |

### 3.6: APTC-PA18 combined with ciprofloxacin has a synergistic action against planktonic *P. aeruginosa*

CIs APTC-DFI-2, APTC-DFI-3 and APTC-DFI-4, all sensitive to APTC-PA18 and to ciprofloxacin, were selected to further investigate the potential for phage and ciprofloxacin synergistic action. APTC-DFI-2 showed a significant decrease in planktonic cells after treatment with ciprofloxacin alone and APTC-PA18 alone in comparison to the untreated control. APTC-PA18 combined with ciprofloxacin further reduced bacterial viability with a significant reduction in comparison to either ciprofloxacin or APTC-PA18 treatment alone. A similar result was observed for the APTC-DFI-3 and APTC-DFI-4. Results are shown in Figure 5.

**Figure 5.**
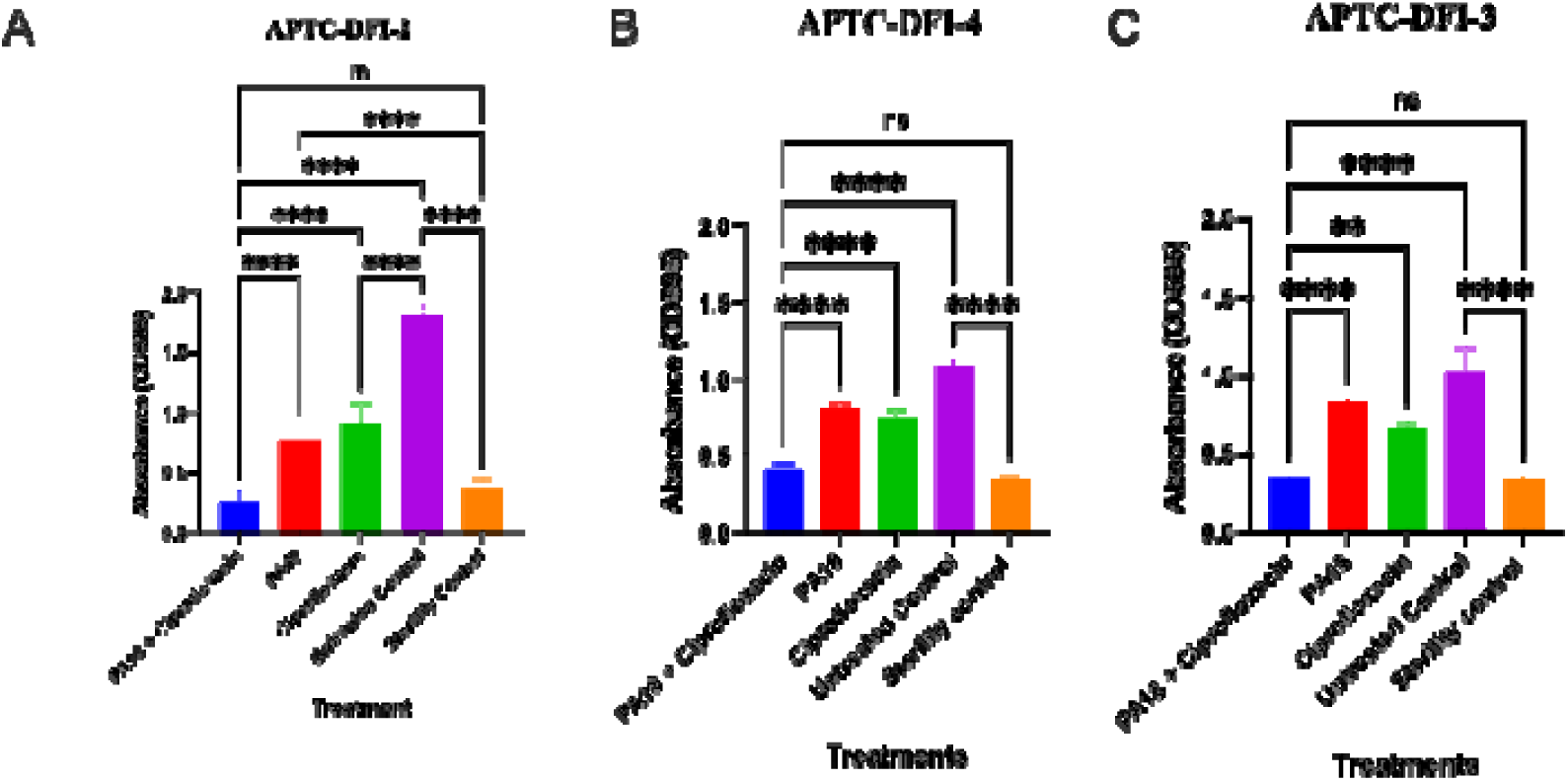
Phage APTC-PA18 and ciprofloxacin combination treatment. The planktonic cells of the CIs in graphs (A) APTC-DFI-2, (B) APTC-DFI-4 and (C) APTC-DFI-3 were treated with APTC-PA18 (MOI = 1) combined with ciprofloxacin (1/2 MIC) (blue bar), APTC-PA18 alone (10^8^ PFU/mL) (red bar) and ciprofloxacin alone (1/2 MIC) (green bar). An untreated control (purple bar) and a sterility control (orange bar) with fresh media were included. OD595 was measured 24 hours later. Data expressed as mean ± SEM for three independent experiments. ns = not significant; ^**^ = p<0.01; ^****^ = p <0.0001. PA18= APTC-PA18.

### 3.7: APTC-PA18 combined with ciprofloxacin has synergistic anti-biofilm effects

We then tested the potential for synergistic antibiofilm effects using the combination of phage with ciprofloxacin. Ciprofloxacin at 1 MIC (for APTC-DFI-2), 2 MIC (for APTC-DFI-4) and 3 MIC (for APTC-DFI-3) did not significantly reduce the bacterial biomass compared to the control after 24 hours. APTC-PA18 at 10^8^ PFU/mL alone did not reduce the biomass of APTC-DFI-2 but the biomass of APTC-DFI-4 and APTC-DFI-3 was significantly reduced compared to the untreated control (p=0.0019 and p=0.0069 respectively). The APTC-DFI-2 showed a significant synergistic reduction in biofilm biomass when treated with APTC-PA18 at 10^8^ PFU/mL in combination with ciprofloxacin in comparison to similar concentrations of APTC-PA18 and ciprofloxacin treatments alone. Similar results were obtained for APTC-DFI-4 and APTC-DFI-3. Results are shown in Figure 6.

**Figure 6:**
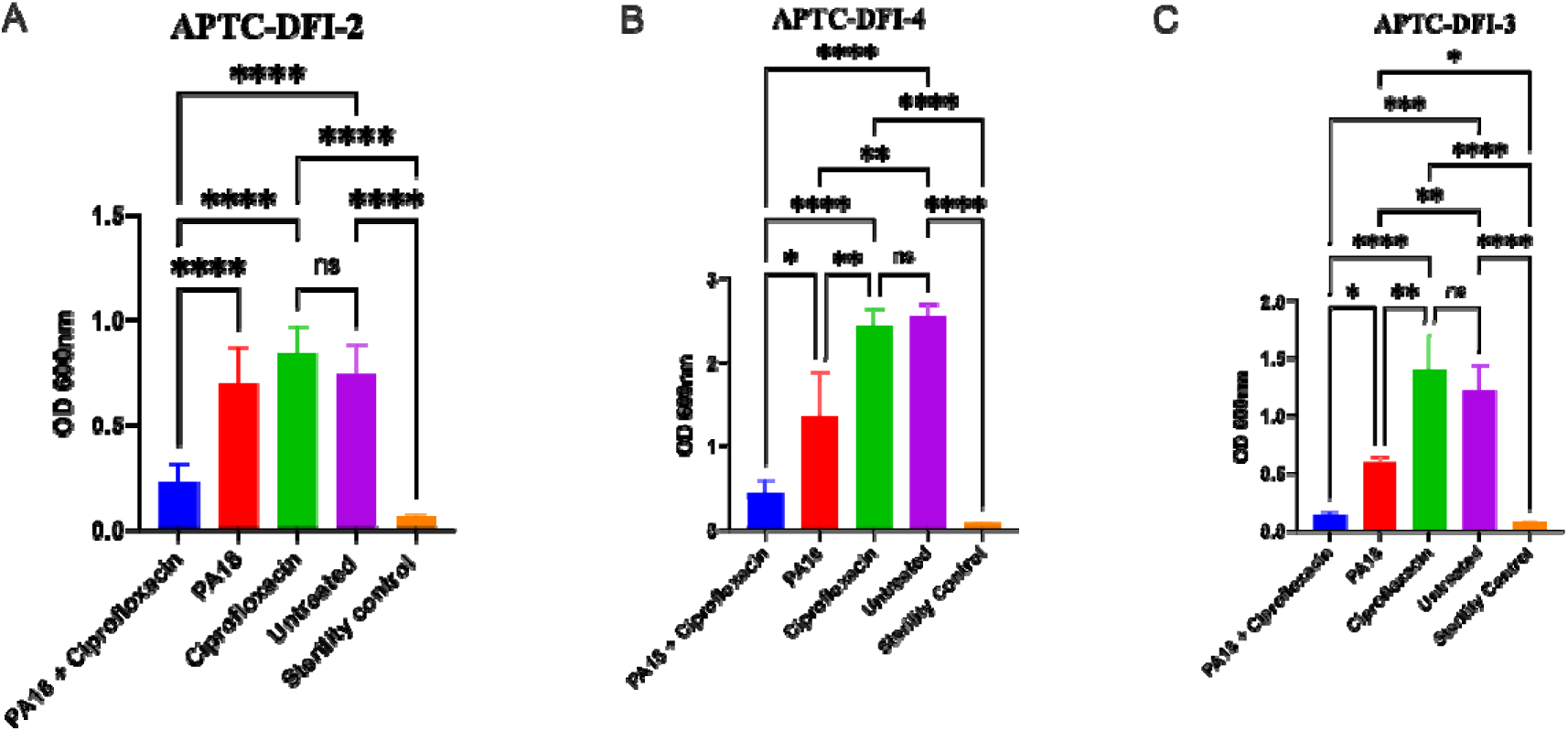
Anti-biofilm effects of the combination of APTC-PA18 and ciprofloxacin. The biofilms of the CIs (A) APTC-DFI-2, (B) APTC-DFI-4 and (C) APTC-DFI-3 were grown for 24 hours and treated with APTC-PA18 (10^8^ PFU/mL) combined with ciprofloxacin (blue bar), APTC-PA18 alone (10^8^ PFU/mL) (red bar) and ciprofloxacin alone (green bar). An untreated control (purple bar) and a sterility control (orange bar with fresh media) were included. OD_600_ was then measured 24 hours post-treatment to determine biofilm biomass. Data expressed as mean ± SEM for three independent experiments. ns = not significant; ^*^ = p<0.05; ^**^ = p<0.01; ^***^; p<0.001; ^****^ = p<0.0001.

**Figure 7:**
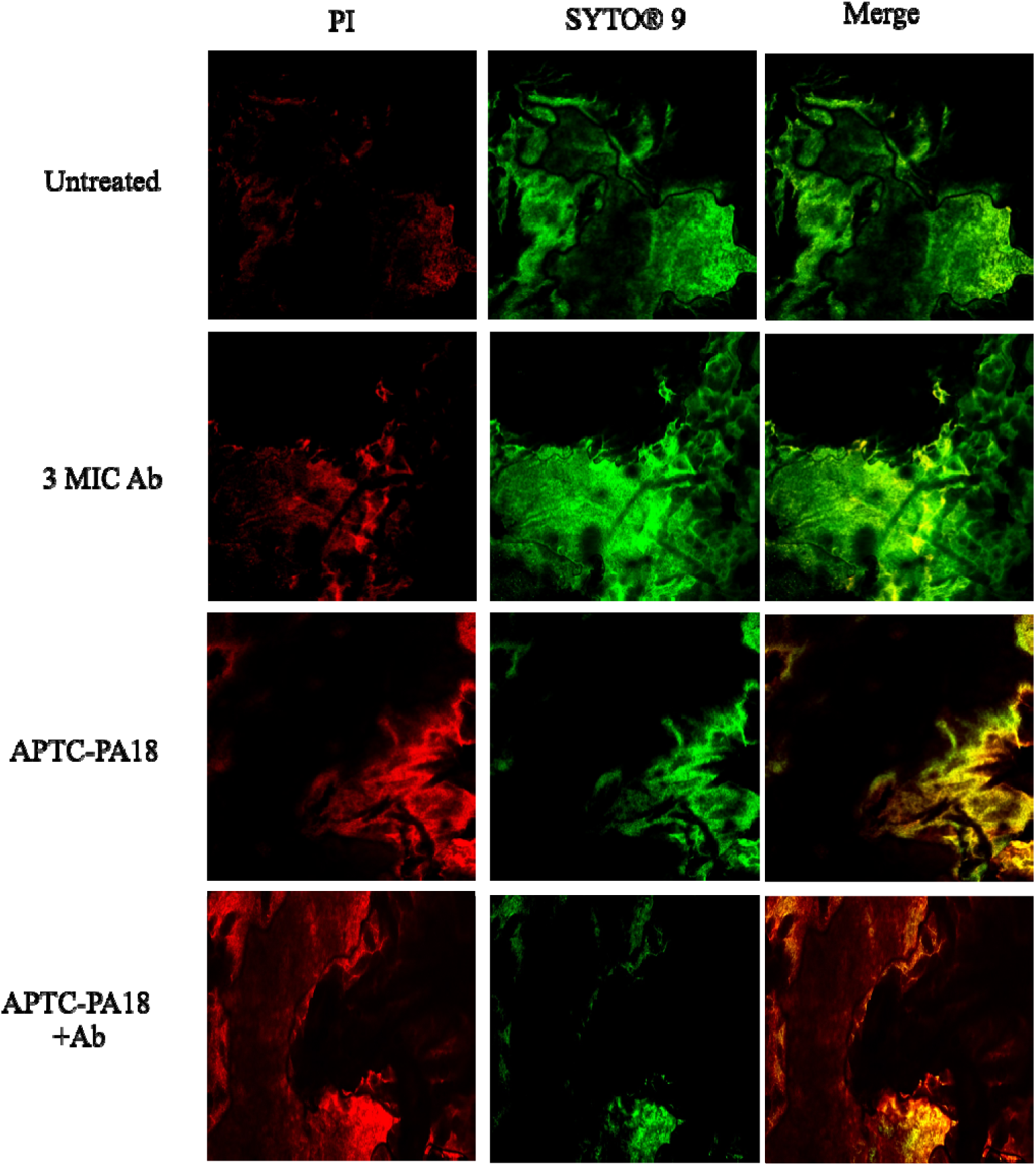
Phage and ciprofloxacin synergistic *ex vivo* treatment. Fresh tissue with *P. aeruginosa-positive* infection was harvested and treated with phage APTC-PA18, 3 MIC ciprofloxacin and APTC-PA18-ciprofloxacin combination for 24 hours. LIVE/DEAD^®^ Bacterial Viability Kit was used in showing dead (PI, red) and live (SYTO® 9, green) biofilm. Ab =ciprofloxacin.

### 3.8: APTC-PA18 in combination with ciprofloxacin reduces the biofilm biomass of DFU tissue *ex vivo*

To confirm the synergistic anti-biofilm effects of the APTC-PA18 phage and ciprofloxacin combination treatment, DFU *P. aeruginosa* (APTC-DFI-8) infected tissue was harvested and treated for 24 hours with phage, ciprofloxacin, or phage-ciprofloxacin in combination. The untreated control showed evidence of viable *P. aeruginosa* biofilm. In comparison to the untreated control, treatment with 3 MIC ciprofloxacin had minimal effects on reducing the biofilm biomass. APTC-PA18 phage treatment alone reduced the biofilm viability. Ciprofloxacin treatment in combination with phage appeared to have the strongest effect with evidence of bacterial cell death throughout the sample.

## 4: Discussion

*P. aeruginosa* is one of the most prevalent pathogens cultured from patients with diabetes-related foot infection (DFI), according to our study. This was further confirmed by 16S rRNA long read sequencing where *P. aeruginosa* was also shown to be among the top 3 pathogens with the highest relative abundance. Bacteriophage PA18 was isolated from DFI and shown to be lytic with a broad host range infecting 7/8 *P. aeruginosa* CIs from DFI. The combination of APTC-PA18 with ciprofloxacin had synergistic effects against *P. aeruginosa* in both planktonic and biofilm forms.

DFI tend to be polymicrobial in nature, despite the microbiota of DFUs being less diverse than that of healthy skin ^29, 30^. Understanding which pathogens dominate the microbiota of DFUs is essential when developing a phage cocktail, as it underlies the ability of the cocktail to target the correct bacteria and resolve infections ^31^. Ciprofloxacin was chosen as a candidate antibiotic for our study as it is commonly used in the management of *P. aeruginosa* infections of diabetes-related foot ulcers (DFUs) and has been shown to have a synergistic anti-microbial effect when used in combination with phage both *in vitro* and *in vivo* ^9, 10^. However, in the clinical context, *P. aeruginosa* in DFUs can become resistant to ciprofloxacin, in particular as a result of prolonged antibiotic treatment ^32^. This resistance is detrimental to infection management and leaves few options for patients, highlighting the need for phage therapy to assist in addressing resistant strains.

The *P. aeruginosa* phage APTC-PA18 is a lytic phage isolated from a DFU *P. aeruginosa* infection, which is a common occurrence, as lytic phages are frequently isolated from active infection sites in humans ^19^. APTC-PA18 was shown to have desirable properties that make it an excellent clinical phage candidate as it lacked lysogeny-associated and toxin-encoding genes. APTC-PA18 is stabled at a wide range of pH and temperatures. Considering the pH of wound fluid of infected DFUs is between pH 6.2 and 8.5 ^33^, and temperature can affect phage attachment, penetration and multiplication and is the most important factor affecting activity ^34^. APTC-PA18 is ideal for clinical use. APTC-PA18 also has a wide host range with 85% of *P. aeruginosa* CIs sensitive to the phage and a strong infective efficacy. Taken together, these factors point to APTC-PA18 being a strong candidate phage for therapeutic use.

When *P. aeruginosa* planktonic cells were treated with sub-inhibitory levels of ciprofloxacin in combination with phage APTC-PA18 Synergistic killing was observed. Phage replication inhibition has been shown to occur when using inhibitory levels of antibiotics simultaneously with phage, hence sub-inhibitory levels of ciprofloxacin were used for this study ^27, 35^. The APTC-DFI-2 showed the largest OD decrease in comparison to the single treatments, and this result was confirmed by the spot plate assay which determined it was the closest to complete eradication of planktonic *P. aeruginosa*. The synergistic effect seen in this study may be not only be pharmacodynamic but also evolutionary ^8^. Namely, mutations in genes encoding for efflux pumps and the genes gyrAB and parCE, allow *P. aeruginosa* to upregulate efflux pumps and reduce the affinity DNA gyrase of Topoisomerase IV to ciprofloxacin as a means of resistance ^36^. Phages in the *Myoviridae* family can bind to surface proteins on drug efflux pumps which exerts pressure on *P. aeruginosa* to alter these efflux pumps in a bid to combat phage infection and develop phage resistance. However, a genetic trade-off occurs wherein the efflux pump’s ability to extrude ciprofloxacin becomes impaired and this increases the bacteria’s susceptibility to ciprofloxacin ^8^. It is undetermined whether increased phage activity, improved effect of ciprofloxacin, or both, are responsible for the synergistic effect in this study.

Biofilms are a hallmark of the chronicity of DFUs and enable infections to be increasingly refractory to antibiotic treatments ^6, 37^. In this study, the biofilms of all 3 CIs tested did not respond to treatment with ciprofloxacin alone, even when concentrations as high as 3 times MIC values were used. In contrast, two CIs (APTC-DFI-3 and APTC-DFI-4) showed a significant reduction in biofilm biomass when treated with APTC-PA18 alone, indicating good antibiofilm activity of phage, as shown in previous studies ^19, 38, 39^. A synergistic reduction in biofilm biomass of all 3 *P. aeruginosa* CI biofilms grown *in vitro* and treated with the APTC-PA18-ciprofloxacin combination was seen and the combination treatment was also most effective against *P. aeruginosa* biofilm infection of DFU tissue. This synergistic anti-biofilm effect could potentially be explained by better biofilm penetration by APTC-PA18 and ciprofloxacin. Studies have shown that phage uses depolymerase to degrade polysaccharides in the biofilm and penetrate it as well as infect persister cells within the biofilm ^9^. Whilst no depolymerase genes were identified in the APTC-PA18 genome, some genes were uncharacterised and thus, without further investigation, it cannot be excluded that some of those genes might encode for proteins with depolymerase activity.

In conclusion, this study identified *P. aeruginosa* amongst the most common and abundant pathogens in infected DFUs in our patient cohort and that such infected sites can harbour lytic phages that can be isolated and developed as a therapy. The isolated novel *P. aeruginosa* specific phage, APTC-PA18 was shown to have many desirable properties that make it a good therapeutic candidate against *P. aeruginosa* infections and biofilms. The synergistic anti-microbial effect of APTC-PA18 in combination with ciprofloxacin, a clinically relevant antibiotic, was exhibited against both planktonic and biofilm forms of *P. aeruginosa* CIs *in vitro* and *ex vivo*. These findings are of clinical significance, as this combination therapy could be an effective approach in managing *P. aeruginosa* infections of DFUs.

## Declarations

### Ethical Approval

HREC/14/TQEH/603.

### Sequence Information

APTC-PA18: GenBank OP957382; 16s sequencing: SAMN32081788 to SAMN32081812 (bioproject: PRJNA909342).

